# A multistate capture-recapture model to estimate cause-specific injury and mortality of North Atlantic right whales

**DOI:** 10.1101/2023.10.15.562416

**Authors:** Daniel W. Linden, Jeffrey A. Hostetler, Richard M. Pace, Lance P. Garrison, Amy R. Knowlton, Véronique Lesage, Rob Williams, Michael C. Runge

## Abstract

Understanding the causes of mortality for a declining species is essential for developing effective conservation and management strategies, particularly when anthropogenic activities are the primary threat. Using a competing hazards framework allows for robust estimation of the cause-specific variation that may exist across multiple dimensions, such as time and individual. Here, we estimated cause-specific rates of severe injury and mortality for North Atlantic right whales (*Eubalaena glacialis*), a critically endangered species that is currently in peril due to human-caused interactions. We developed a multistate capture-recapture model that leveraged 30 years of intensive survey effort yielding sightings of individuals with injury assessments and necropsies of carcass recoveries. We examined variation in the hazard rates of severe injury and mortality due to entanglements in fishing gear and vessel strikes as explained by year and the age and reproductive status of the individual. We found strong evidence for increased rates of severe entanglement injuries after 2013 and for females with calves, with consequently higher marginal mortality. The model results also suggested that despite vessel strikes causing a lower average rate of severe injuries, the higher mortality rate conditional on injury results in significant total mortality risk, particularly for females resting from a recent calving event. Large uncertainty in the estimation of carcass recovery rate for vessel strike deaths permeated into the apportionment of mortality causes. The increased rates of North Atlantic right whale mortality in the last decade, particularly for reproducing females, puts the species at risk of severe decline. By apportioning the human-caused threats using a quantitative approach with estimation of relevant uncertainty, this work can guide development of conservation and management strategies to facilitate species recovery.

## 1. Introduction

Identifying the mechanisms responsible for demographic change is essential for effective conservation and management of animal populations (Williams et al., 2002), particularly for species that are heavily influenced by human activity. Human-caused mortality due to hunting or harvest effort may be modified with relatively straightforward strategies (Nichols et al., 2007; Koons et al., 2014), while deaths that result as a byproduct of other anthropogenic activities generally present a greater challenge for mitigation (Van Der Hoop et al., 2013; May et al., 2015; Benson et al., 2023). In either case, robust approaches to estimating the cause-specific mortality rates experienced by wild populations can assist with designing management strategies that effectively target the relevant mechanisms (Heisey and Patterson, 2006; Tavecchia et al., 2012; Nater et al., 2020).

Observations of mortality can take many forms, but typically such records are a small sample of total population deaths with variable contamination by imperfect information depending on the system and monitoring scheme. Large-scale banding and tagging studies can yield sample sizes suffiicient for statistical modeling, especially for species that are harvested (Brownie, 1978; Lebreton et al., 1992), but in some systems the fraction of total mortality represented by carcass detections is potentially too low for useful inferences (e.g., Williams et al., 2011). Opportunistic carcass encounters of non-hunting related mortality events, typical in marine systems (e.g., strandings), can result in variable carcass conditions that degrade the ability to determine cause of death (Moore et al., 2020). Even for telemetered individuals with negligible detection concerns, cause-of-death determination can be equivocal (Cristescu et al., 2022). Regardless of sampling approach, proper estimation of cause-specific mortality rates will often need to accommodate multiple levels of observational uncertainty.

North Atlantic right whales (*Eubalaena glacialis*; hereafter, NARWs) are a critically endangered species with multiple anthropogenic mortality causes that are currently responsible for its plight, primarily entanglement in fishing gear and vessel strike (Kraus and Rolland, 2009; Knowlton et al., 2022). An intensive monitoring program yields high sighting rates with >80% of individuals typically observed each year (Pace et al., 2017), yet uncertainty remains regarding the degrees to which entanglements and vessel strikes each contribute to total deaths with ∼2/3 of mortality events being unobserved (Pace et al., 2021). The observed carcasses with successful necropsy have been used to make inferences about the various mortality threats to NARWs (Sharp et al., 2019; Henry et al., 2021) while recognizing that the cause apportionment within the sample may not be entirely representative due to biases resulting from variation in the persistence and detection of certain deaths (Moore et al., 2020). Evidence suggests a higher prevalence of entanglement-related deaths given the high rates of entanglement injury and subsequently poor individual health (Schick et al., 2013; Rolland et al., 2016; Knowlton et al., 2022) and the apparent disappearance of severely entangled individuals from the sightings database (Pace et al., 2021). Rapid population decline of NARWs during the last decade (Hayes et al., 2022) has emphasized the importance of partitioning mortality causes to guide management strategies.

In this paper we developed a multistate capture-recapture model (Lebreton et al., 2009) to estimate cause-specific rates of transition between states of severe injury and death for NARWs. We posited that severe injuries observed in live individuals could help indicate the cause of subsequent mortality events, allowing us to leverage the immense monitoring efforts (Hamilton et al., 2007) and health assessments (Knowlton et al., 2016; Pettis et al., 2017) that exist for the species. These monitoring data have been used to model and predict fine-scale changes in NARW health due to multiple stressors (e.g., Knowlton et al. (2022); Pirotta et al. (2023); and references therein). Here, we integrated 30 years (1990–2019) of sightings data with carcass recovery and necropsy information (Moore et al., 2004; Sharp et al., 2019) and explicitly modeled the observation processes to improve inferences on mortality rate estimation. Crucially, we allowed for state uncertainty (i.e., “multievent” modeling; Pradel (2005)) to accommodate injuries and deaths without clear causes. We explored injury and mortality variation across individual life-history stages with an emphasis on breeding females, in addition to changes in rates across time. Our modeling framework enables a partitioning of mortality causes with uncertainty to provide context for how anthropogenic threats have influenced NARW population decline in recent years.

## 2. Materials and methods

### 2.1. Monitoring data

Monitoring efforts by the North Atlantic Right Whale Consortium (https://www.narwc.org/) have covered most of the species range in the western Atlantic since 1980 (Kraus and Rolland, 2007), though consistent effort began closer to 1990 (Pace et al., 2017). NARWs are individually identifiable due to distinct natural markings (i.e., callosity patterns) that become permanent at a young age (>0.5-years old), and the catalog curated by the New England Aquarium includes high quality photographs that allow for constructing individual sightings histories (Hamilton et al., 2007). In addition to identifying individuals in photographs, a health assessment is conducted to score the severity of observed injuries, among other attributes (Knowlton et al., 2016; Pettis et al., 2017). Data on recovered carcasses of large whales are gathered and maintained by multiple stranding networks situated along the Atlantic coasts of the United States, (*Greater Atlantic Marine Mammal Stranding Network* | *NOAA Fisheries*^3^) and Canada (*Report a marine mammal or sea turtle incident or sighting*^4^). We relied on a detailed list of documented right whale mortalities aggregated from those networks at the Northeast Fisheries Science Center (e.g., Henry et al. (2021)).

We used sightings data extracted on 23 December 2021, which included 691 whales >0.5-years old known to be alive sometime during 1 April 1990–30 September 2019. The data were supplemented by New England Aquarium (A. Knowlton, *unpublished data*) with detected wounding events from inspection of images associated with all sightings of identified individuals when the suite of images was adequate for evaluation. Although multiple levels of wound classification exist (Knowlton et al., 2016), we only considered severe wounds for designating the injury state of an individual. We constructed encounter histories by individual and year with a defined encounter period of 1 April–30 September (i.e., “summer” surveys), during which observations of individual states (or events) could occur. The recorded event *y*_*i,t*_ for an individual *i* in year *t* could take one of eight values: 1 = seen alive; 2 = seen injured by entanglement; 3 = recovered dead by entanglement; 4 = seen injured by vessel strike; 5 = recovered dead by vessel strike; 6 = seen injured by unknown cause; 7 = recovered dead by unknown cause; and 8 = not seen. The value of an event for individuals with multiple sightings during an encounter period was assigned as the most consequential event: mortality > injury > no injury; and vessel strike > entanglement.

In addition to the defined encounter period, we used sightings collected from the southern Atlantic coast during the previous calving season (generally 1 Dec–30 Mar; “winter” surveys) to provide evidence of reproductive state as a year-specific individual attribute. The winter surveys are believed to provide a near-complete census of calf production. As such, individuals were classified as “not with calf,” “with calf,” and “recently with calf” (i.e., with a calf during the prior year) to reflect reproductive states that were hypothesized to influence injury and mortality rates. Other individual attributes included age class and sex which were known for 85% and 94% of individuals, respectively.

### 2.2. A multistate model for observed injury and mortality events

#### 2.2.1. State transition matrix

Our multistate capture-recapture model described transitions between individual states to estimate rates of severe injury and death (Figure 1). We defined the following true states: 1 = alive; 2 = injured by entanglement; 3 = recent death by entanglement; 4 = injured by vessel strike; 5 = recent death by vessel strike; and 6 = dead. Death from a particular cause was conditional on obtaining a severe injury from said cause. For example, an individual *i* that is alive and uninjured in year *t* can obtain a severe injury due to entanglement with probability 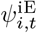 and, if having obtained a severe entanglement injury, survive to the next year with probability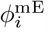. Under this model formulation, mortality from causes other than anthropogenic injuries is not possible.

**Figure 1:**
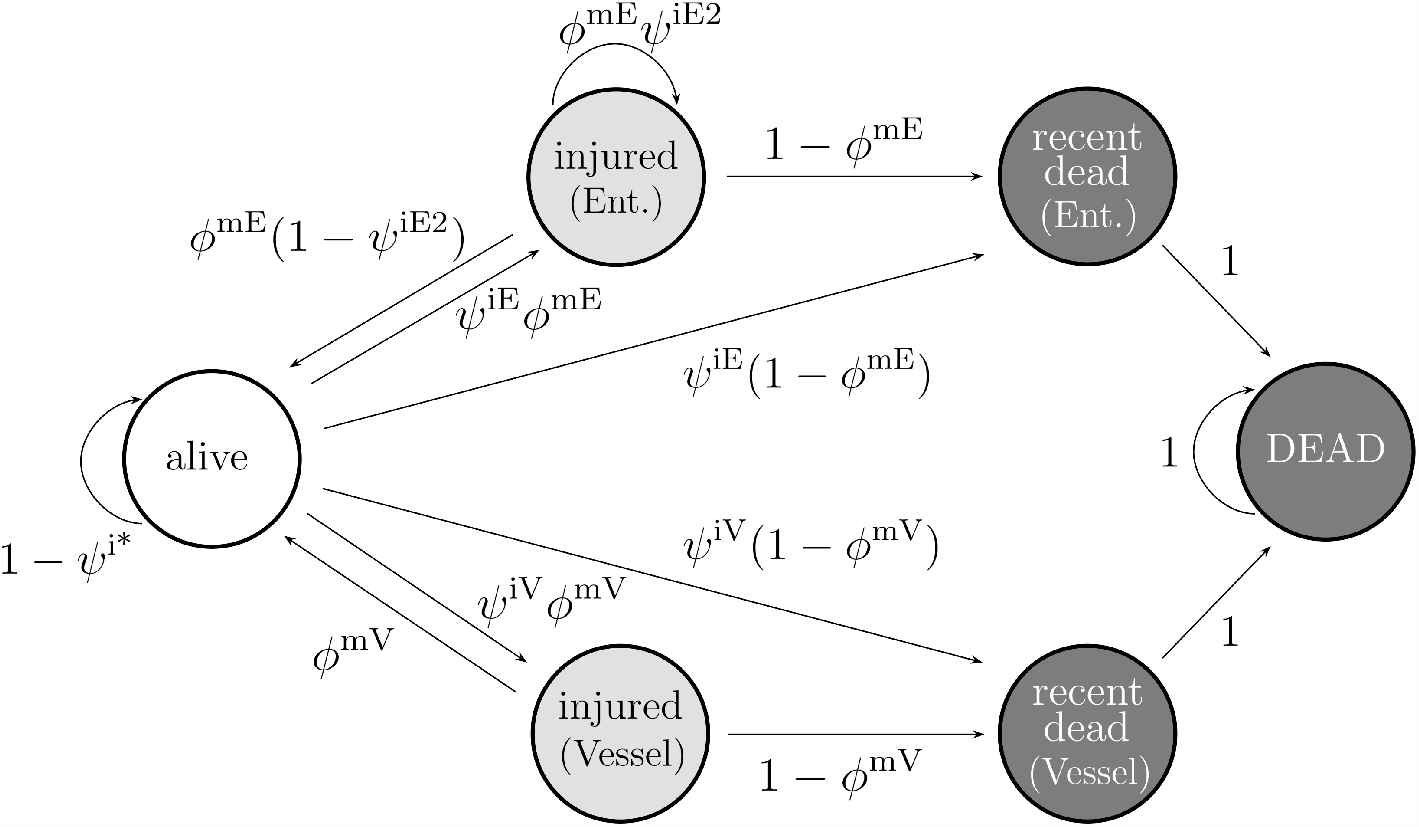
State transition model for North Atlantic right whale injury and mortality states across annual time steps. Probabilities include those for obtaining an injury due to entanglement (*ψ*^iE^) or vessel strike (*ψ*^iV^), keeping an entanglement injury (*ψ*^iE2^), obtaining an injury due to any cause (*ψ*^i*^), and surviving an injury due to entanglement (*ϕ*^mE^) or vessel strike (*ϕ*^mV^). Subscripts for indexing individual *i* and year *t* are omitted for clarity.

We assumed that natural mortality was negligible for two reasons. First, non-calf NARWs have never been observed to die from natural causes according to necropsy evidence (Moore et al., 2004; Sharp et al., 2019). Most whales with an unidentified cause of death exhibit evidence of injury consistent with one or more anthropogenic causes. Second, metabolic theory indicates that natural mortality must necessarily be negligible for a majority of a species lifespan, until senescence at the very end (Anderson, 2018). Using the closely related bowhead whale (*Balaena mysticetus*) as an approximation, with a lifespan of >200 years (Mayne et al., 2019), we therefore posited that natural mortality has not played a significant role, calf recruitment aside, in the recent history of NARW population dynamics.

Probabilities of injury and mortality were allowed to vary by individual attributes and year according to the following state transition matrix:

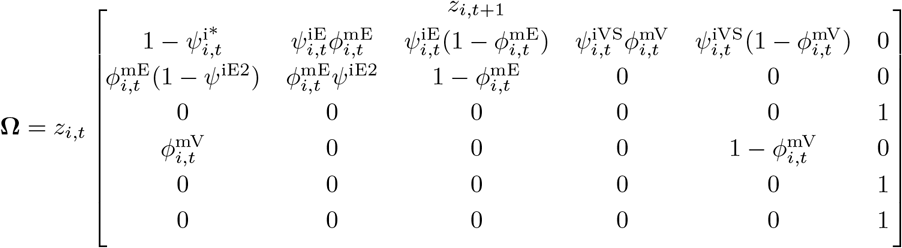

The state transition matrix makes clear that the only path to a recently dead state is through injury. Individuals that avoid severe injury, according to the inverse probability of obtaining any injury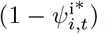, automatically transition back to state 1 in the following year. In addition to obtaining a new injury, we defined *ψ*^iE2^ as the probability of retaining a severe entanglement injury for an additional year, possibly due to attached gear. We assumed that severe vessel injuries either heal enough the following year to no longer be classified as severe (hence, *z*_*t*_ = 4 transitions to *z*_*t*+1_ = 1) or cause a recent death (*z*_*t*+1_ = 5).

The severe injury causes represented competing risks, so to properly accommodate individual and temporal variation we parameterized the model using hazard rates (Ergon et al., 2018; Nater et al., 2020). The probability of obtaining any injury was therefore a function of the combined hazard rates of each cause:

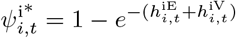

where 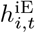 and 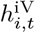 are the hazard rates for injury due to entanglement and vessel strike, respectively. Given these hazard rates, the unconditional probabilities of injury as specified in the transition matrix can be calculated as follows:

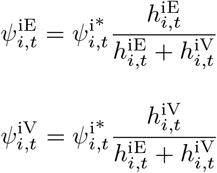

We also defined the mortality probabilities in terms of hazard rates such that 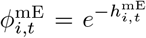 and 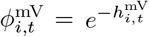. Since mortality was conditional on injury and, thus, the risks of mortality were not competing, the hazard formulation simply amounted to using a loglog link for modeling variation in the probabilities (Ergon et al., 2018).

We hypothesized multiple sources of potential variation for the hazard rates across individuals and years. Previous evidence suggests younger animals and females recovering from birthing and calving activities are more vulnerable to mortality (Knowlton et al., 2022). Additionally, NARW distribution appeared to shift sometime after 2010, when large-scale ecosystem changes in the western Atlantic resulted in altered prey availability and increased use of Canadian waters (Davis et al., 2017; Meyer-Gutbrod et al., 2018; Davies and Brillant, 2019). As such, we used the following general structure for the log-linear models of injury and mortality hazard rates:

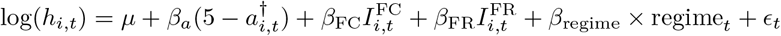

Here, *h*_*i,t*_ is the hazard rate for injuries due to cause 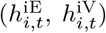 or mortalities due to cause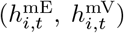. Each of the coefficients likewise has four versions, corresponding to the two injury rates and the two mortality rates; *μ* is the log-scale baseline hazard rate; *β*_*a*_ is the effect of early age on the hazard; 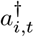 is the age of the whale if that age is <5, and 5 otherwise; *β*_FC_ is the effect of being a female with a calf; 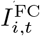 is an indicator variable that is 1 if individual *i* is a female with a calf in year *t* and 0 otherwise; *β*_FR_ is the effect of being a resting female; 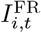 is an indicator variable for resting female; *β*_regime_ is the effect of a regime shift in 2013; regime_*t*_ is a contrast variable taking values of -1 (years up to 2013) or 1 (years following 2013) for each year *t*; and *ϵ*_*t*_ is the random normal effect of year *t* on the hazard rate. The regime definition was chosen as a compromise between 2010 and 2015, a period that had resulted in fewer sightings from traditional survey locations due to an apparent shift in the whale distribution. Initial model fitting suggested little evidence of annual variation in the two mortality hazard rates so the regime effects and temporal random effects were removed from consideration and indexing for year *t* was unnecessary (i.e., 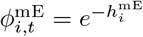 and 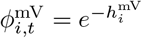). We constrained the probability of retaining an entanglement injury (*ψ*^iE2^) to be constant across individuals and time.

#### 2.2.2. Observation matrix

The observations *y*_*i,t*_ were conditional on true states *z*_*i,t*_ as dictated by the observation matrix (**Θ**), which linked each true state to the possible observations that could occur in a given year. Sighting (*p*_*i,t*_) and recovery 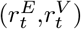 probabilities both varied by year, with the former also varying by individual and the latter by cause of death. The probabilities of identifying cause of severe injury (*ρ*^*inj*^) and mortality (*ρ*^*mort*^) were constant and simply apportioned the unknown injuries and deaths between the two known causes. The observation matrix was thus defined as follows:

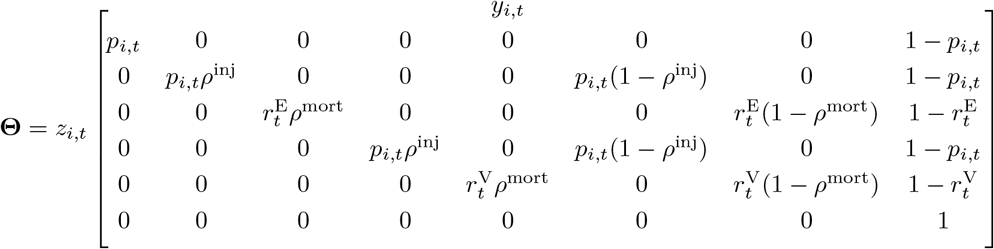

We modeled variation in sighting probabilities and recovery rates using logit-linear models:

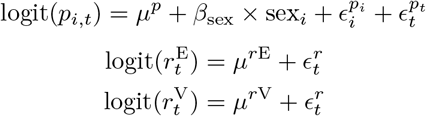

Here, *μ*^*p*^ is the logit-scale mean sighting probability; *β*_sex_ is the effect of the contrast variable sex_*i*_ (−1 = male, 1 = female); 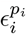 is the random normal effect of individual *i*; and 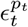 is the random normal effect of year *t*. The recovery probabilities were likewise specified to have separate logit-scale means (*μ*^*r*E^, *μ*^*r*V^), though they shared a single random normal effect 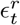 for each year.

### 2.3. Model fitting and assessment

We fit the model in R (R Core Team, 2022) using Markov chain Monte Carlo (MCMC) with NIMBLE (de Valpine et al., 2017, 2022) and a dynamic hidden Markov model likelihood (Turek et al., 2016; Goldstein et al., 2021). This likelihood marginalizes the latent states for increased computational efficiency (Turek et al., 2016) while still leveraging the Bayesian capability to robustly estimate random effects across multiple dimensions with MCMC. Although it was not strictly necessary, we conditioned on first capture (f_*i*_) and did not estimate distinct initial state probabilities. The general likelihood for the sightings data was therefore defined as follows (see Goldstein et al. (2021) for more details):

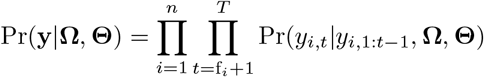

The likelihood was also conditional on individual attributes, and imputation was necessary for some missing values. Individuals with an unknown initial age or sex were assigned values according to probability distributions. The distribution of observed known ages was not informative for unknown ages given that known-age individuals had either been sighted as 0.5-yr-old animals or had ancillary information available (e.g., genetic sample or sighting prior to 1990). Initial ages were therefore drawn from a categorical distribution *a*_*i*,f_ *∼* Cat(***π***) with probabilities that depended on whether age was known. For known ages, *π*_1:5_ *∼* Dir(***α***) with ***α*** = [1, 1, 1, 1, 1]; for unknown ages, *π*_1_ = 0 and *π*_2:5_*∼* Dir(***α***) with ***α*** = [16, 8, 4, 2]. The latter construct assumes unknown age animals were not first seen in their birth year (given the obvious smaller size of 0.5-year olds) but most likely to be younger than adults. Unknown sex animals had values drawn from the observed sex distribution such that *sex*_*i*_*∼* Bern(*δ*), where *δ* indicated the probability of being female.

We assigned vague priors for most parameters, including Gamma(1, 1) for the hazard rate intercepts, Uniform(0, 1) for probabilities, and Normal(0, *σ* = 3) for effects coefficients. We assigned informative priors for the variance parameters of the random error terms where *ϵ∼* Normal(0, *σ*); each *σ* was drawn from a scaled Half Student-T (Rankin et al., 2016) with *τ* = 4 and df = 6 for the hazard rates and recovery probabilities (more constraint) and *τ* = 1 and df = 3 for the sighting probabilities (little constraint). For the cause-specific recovery probabilities, we used a slightly informative Beta(2, 4) prior to ensure a reasonable range of values consistent with previous estimates of carcass recovery (Pace et al., 2021). To improve mixing, a sum-to-zero approach was used for the random effects (Ogle and Barber, 2020).

We used a model selection technique (variable selection by reversible jump MCMC; de Valpine et al. (2022)) that estimated probabilities of inclusion for covariates of individual attributes in the log-linear models of injury and mortality. This resulted in some covariate effects being set to 0 during some MCMC iterations, allowing us to test the age and reproductive state effects for the hazard rates while attempting to avoid over-parameterization.

The MCMC algorithm was run for 25,000 iterations over 2 chains, after a burn-in of 5,000 iterations. We examined trace plots and the potential scale reduction factor (R-hat; Brooks and Gelman (1998)) to assess convergence and calculated the prior-posterior overlap for the main parameters to assess identifiability (Gimenez et al., 2009). Finally, we assessed model fit with posterior predictive checks (PPC) (Conn et al., 2018) to compare annual counts of events observed to those predicted. We followed the strategy used by Nater et al. (2020) of selecting 500 posterior samples and simulating 10 sightings histories for each sample to generate distributions of predicted events for comparison with what was observed. We conditioned on the random effects estimated for the hazard rates and recovery probabilities, but not on those estimated as part of sighting probabilities (i.e., 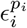 and 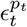); the latter were simulated during the PPC in an attempt to challenge the model with a “mixed predictive check” (Conn et al., 2018).

## 3. Results

The 691 individuals across 30 years yielded the following counts of observations: 7651 not severely injured whales; 75 severe entanglement injuries; 20 entanglement dead recoveries; 4 severe vessel strike injuries; 25 vessel strike dead recoveries; 5 unknown-cause severe injuries; 20 unknown-cause dead recoveries. Summaries of the posterior distributions for model parameters can be found in Table S1, with pairwise scatterplots and correlations between link-scale intercepts for cause-specific injury, mortality, and recovery in Figure S1. Trace plots indicated sufficient convergence with all values of 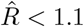. Prior-posterior overlaps were <35% (Table S2) except for the recovery rate of mortalities due to vessel strike (*μ*^rV^), which indicated poor identifiability for this parameter. The posterior predictive check did not indicate any serious lack of fit (Figure S2), though total observed counts of injured animals due to unknown causes (*y*_*i,t*_ = 6) were significantly greater than predicted for 11 of the 29 years assessed. In presenting model results, we report parameter estimates as the posterior median followed with 95% credible intervals in brackets.

### 3.1. Observation processes

Average sighting probability was high with inv-logit(*μ*^*p*^) = 0.794 [0.780, 0.808], consistent with previous mark-recapture estimates of NARWs on an annual time step (Pace et al., 2017), even with our restricting the sightings to spring/summer surveys. Females on average were observed with a lower probability than males 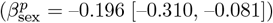 and there was considerable residual variation in the logit-linear model across time 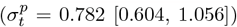 and individuals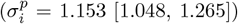. A decrease in average annual sighting probability during the early 2010s is apparent (Figure 2).

**Figure 2:**
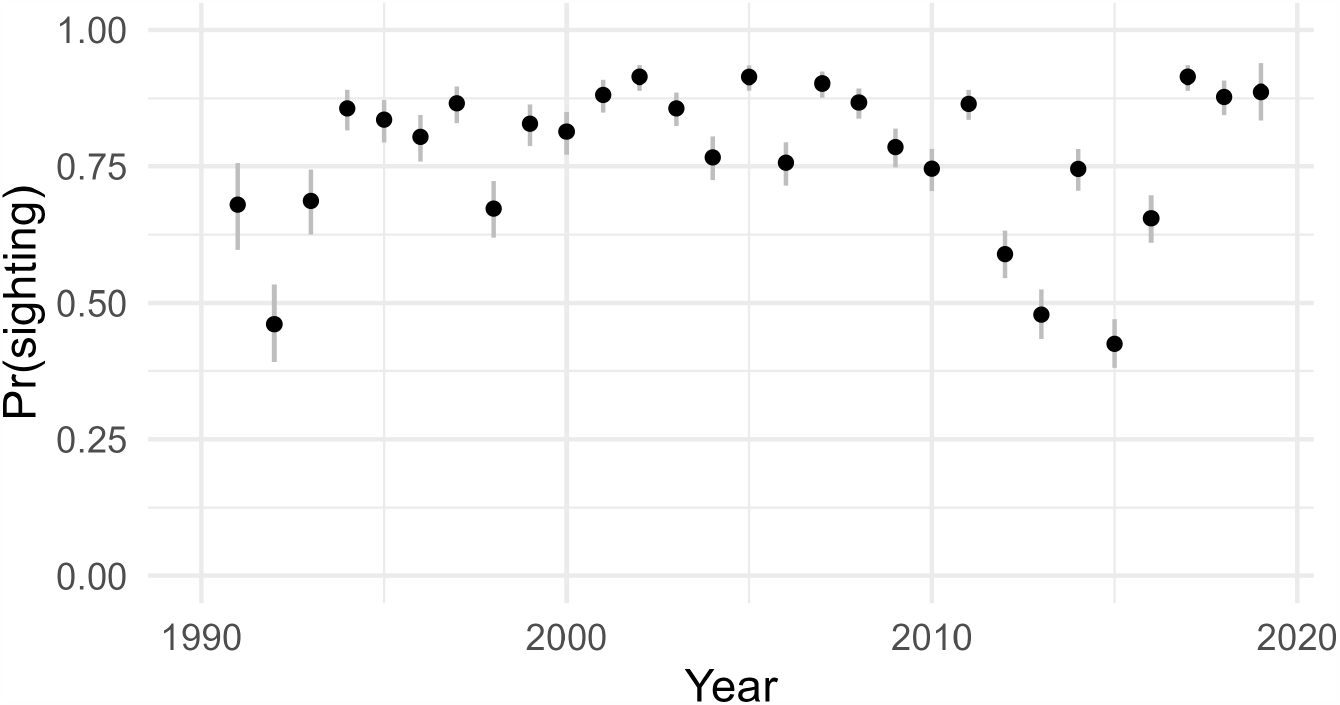
Average annual probability of sighting North Atlantic right whales during spring and summer surveys during 1991-2019.

The probabilities of identifying cause for injuries (*ρ*^inj^ = 0.927 [0.853, 0.971]) and mortalities (*ρ*^mort^ = 0.689 [0.571, 0.792]) both reflected the observed proportions of known vs. unknown causes for each. The carcass recovery rate for entanglement deaths (inv-logit(*μ*^rE^) = 0.129 [0.078, 0.221]) was lower than that for vessel strike deaths (inv-logit(*μ*^rV^) = 0.290 [0.155, 0.601]), though the latter was highly uncertain and may not have been identifiable.

### 3.2. Cause-specific injury and mortality

The average injury hazard rate for entanglement (exp(*μ*^iE^) = 0.028 [0.019, 0.038]) was more than twice the injury hazard rate for vessel strike (exp(*μ*^iV^) = 0.012 [0.005, 0.023]). These low rates resulted in nearly equivalent annual probabilities of obtaining a severe injury, whether due to entanglement (0.028 [0.019, 0.037]) or vessel strike (0.012 [0.005, 0.023]). Conditional on obtaining a severe injury, the average mortality hazard rate was almost 3 times greater for vessel strike (exp(*μ*^mV^) = 2.571 [1.673, 3.683]) than for entanglement (exp(*μ*^mE^) = 0.875 [0.607, 1.113]), resulting in a much lower survival probability for individuals severely injured by vessel strike (0.076 [0.025, 0.188]) than those severely injured by entanglement (0.417 [0.328, 0.545]). The probability of retaining a severe entanglement injury between years (*ψ*^*iE*2^), conditional on survival, was 0.083 [0.020, 0.202].

### 3.3. Variation in injury and mortality

Posterior probabilities of inclusion and effect sizes suggested some evidence of stage-based variation in injury and mortality hazard rates (Figure 3; Table S1), more so for reproductive stages than for younger age classes. The strongest evidence was an increased hazard of severe injury by entanglement for females with a calf 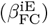 where posterior inclusion was 0.90 and the effect size (1.180 [0.000, 1.750]) was largely above 0. The resulting probability of severe injury for these females increased to 0.090 [0.022, 0.143]. Other stage-based variation had weaker support, though there was some evidence that resting females recently with a calf had an increased hazard of severe injury by vessel strike 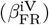given the posterior inclusion (0.66) and effect size (0.795 [0.000, 1.988]).

**Figure 3:**
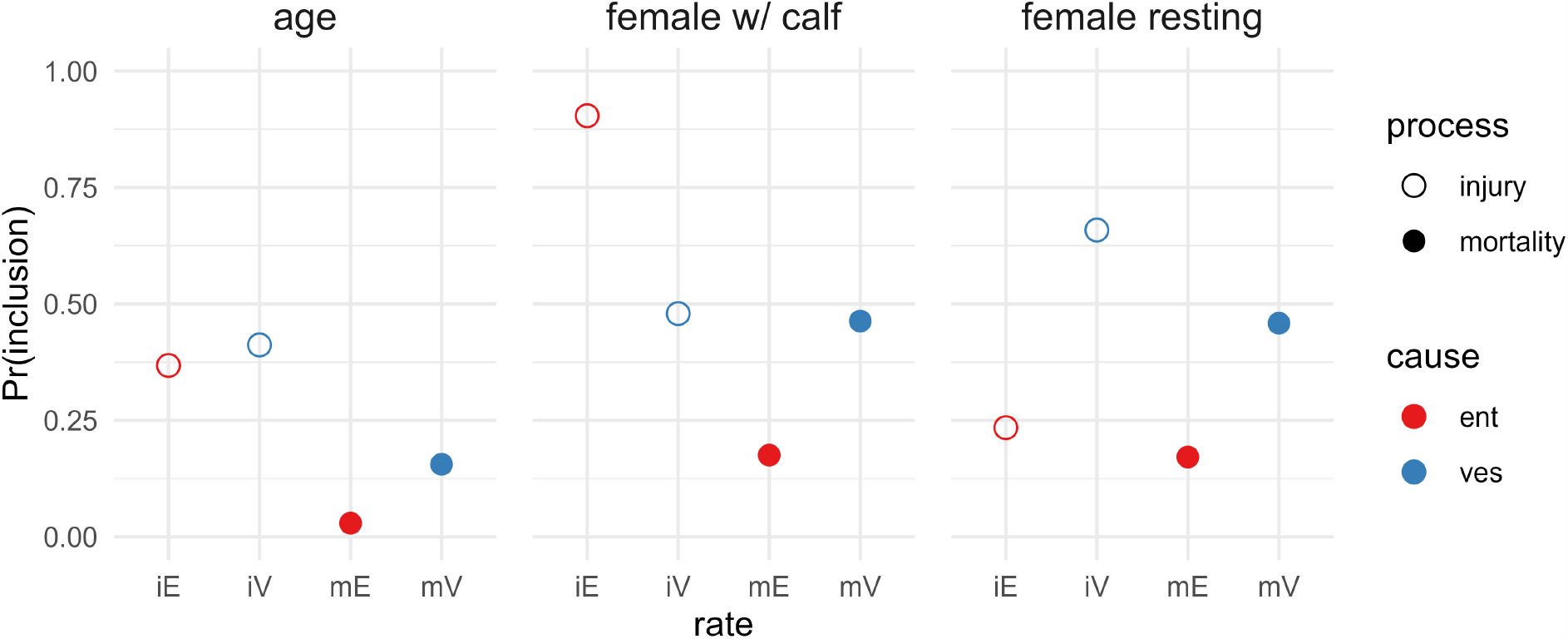
Posterior probability of inclusion for effects explaining variation in injury and mortality hazard rates for entanglement and vessel strike. Coefficients include effects of age, female with a calf (FC), and resting female recently with calf (FR).

Temporal variation in severe injury hazard rates indicated strong support for a regime shift after 2013 in entanglements 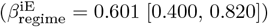 and weaker support for a regime shift in vessel strikes 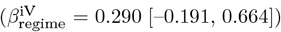. Given that regime_*t*_ was a contrast variable, this resulted in an average probability of severe injury due to entanglement that was much lower prior to the regime shift (0.015 [0.010, 0.022]) than following the regime shift (0.050 [0.033, 0.069]). The weaker evidence of a regime shift in the hazard rate of severe injury due to vessel strike resulted in a smaller difference in the expected probabilities pre-regime shift (0.009 [0.004, 0.016]) and post-regime shift (0.015 [0.005, 0.036]). Residual annual variation in the log-linear hazard rate models was greater for vessel strike injuries (0.370 [0.095, 0.760]) than for entanglement injuries (0.133 [0.009, 0.423]).

The combination of individual and temporal variation in hazard rates resulted in several patterns for the marginal probabilities of NARW mortality (Figure 4). The increased hazard rates of entanglement injury for females with a calf and vessel strike injury for females recently with a calf are both apparent, along with greater uncertainty regarding cause of death. All reproductive states also illustrated the increased injury hazards after 2013 and the subsequently higher marginal probabilities of mortality.

**Figure 4:**
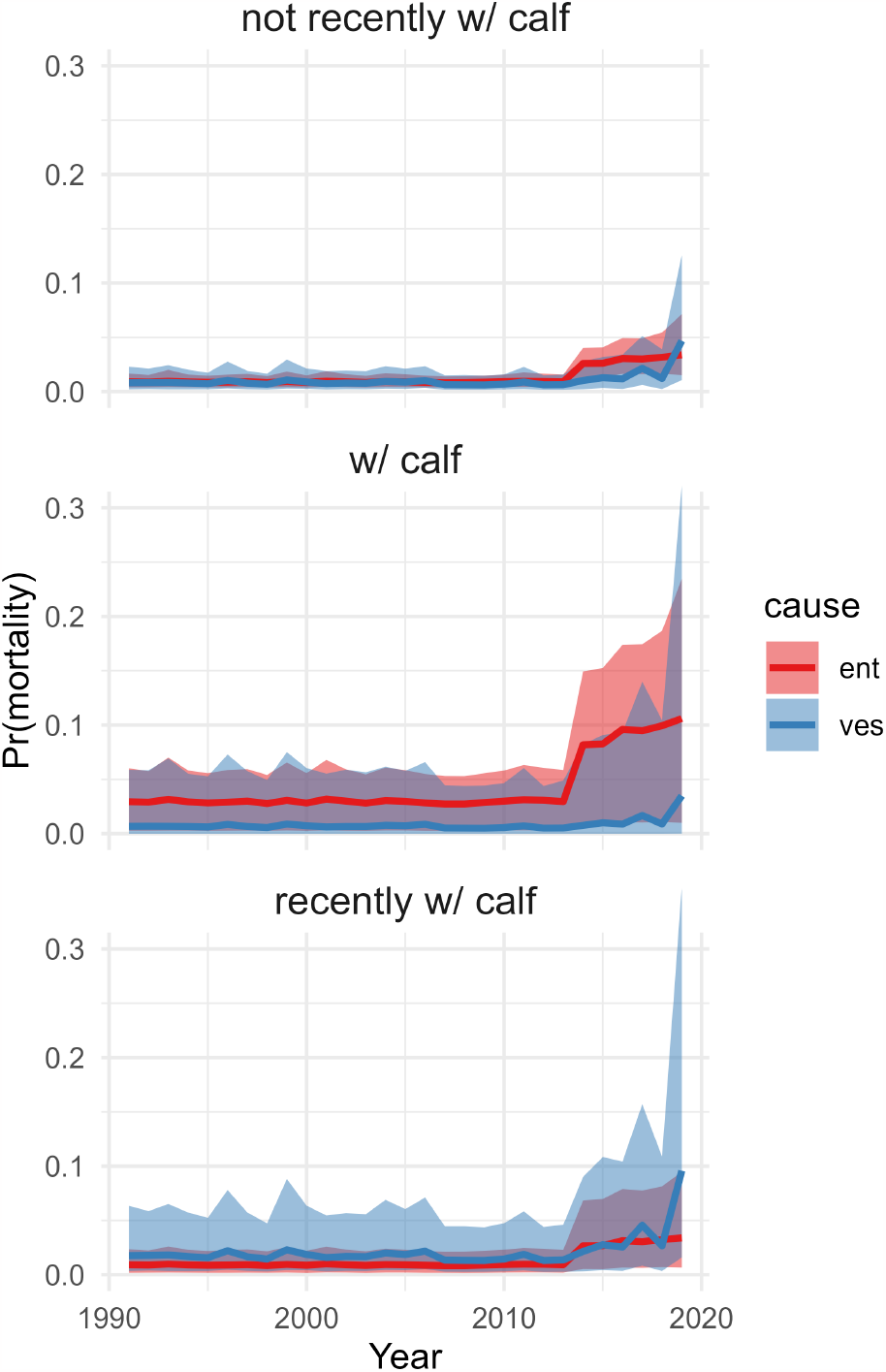
Marginal probability of mortality across time due to both entanglement (red) and vessel strike (blue) for adult whales in one of three reproductive states, including individuals not recently with a calf (males and waiting females), females with a calf (“FC”), and females recently with a calf, i.e., in resting state (“FR”). Median with 95% credible intervals illustrated.

## 4. Discussion

Understanding the causes of mortality for a wild population in decline is a necessary step in designing effective management strategies. In particular, knowledge that mortality risks have changed over time and vary with individual life history stages can facilitate improvements in both monitoring and mitigation. For a critically endangered species like North Atlantic right whales, where deaths primarily occur as a byproduct of human activities, the partitioning of mortality risk is crucial to conservation efforts. We developed a multistate capture-recapture model to leverage the available observations of live and dead whales in a framework that quantifies the hazards and improves understanding of injury and mortality risk to the population. Our construct was similar to an illness-death model (Tassistro et al., 2020), where observations of live individuals are indicative of mortality outcomes. Model results indicated strong evidence for increased rates of severe entanglement injuries after 2013 and for females with calves, with consequently higher marginal mortality. The model also suggested that despite vessel strikes causing a lower average rate of severe injuries, the higher mortality rate conditional on injury results in significant total mortality risk, particularly for females resting from a recent calving event.

The recent plight of NARWs has been well documented by the scientific community thanks to immense efforts by the North Atlantic Right Whale Consortium (NARWC) and its collaborators to monitor the species (e.g., Moore et al., 2021). Stock assessment models using capture-recapture indicate a clear population decline (Pace et al., 2017; Hayes et al., 2022) and individual health assessments with modeling of multiple stressors make clear that anthropogenic threats have been the overwhelming cause of elevated ailments and decreased survival in recent years (Rolland et al., 2016; Knowlton et al., 2022; Pirotta et al., 2023). In addition to direct anthropogenic threats, climate change has altered NARW prey availability and instigated a distribution shift that has likely exacerbated human-caused interactions (Meyer-Gutbrod et al., 2021). Both the complexity of the system and the current state of knowledge about the species are high enough that multiple analytical approaches to understanding NARW population dynamics are warranted. Our attempt here was to provide a framework for modeling the observation processes and quantifying the anthropogenic threats in a manner that could directly inform the development of future population projections and facilitate potential management interventions.

The mechanisms potentially responsible for the estimated variation in the hazard rate of severe entanglement injuries are difficult to tease apart. Females with calves are known to be vulnerable to environmental stressors given the physiological demands of lactation (Rolland et al., 2016), and could therefore incur entanglements at similar rates to other demographic groups but have higher rates of severe injuries resulting from poorer body condition. Conversely, the movement behavior of a female NARW with a calf after birthing, including the migration from the southeast U.S. coast to the northwestern Atlantic and subsequent summer foraging demands, could result in different spatiotemporal exposure to entanglement risk that results in more severe injuries. Additionally, the increased hazard rate of severe entanglement injuries after 2013 is likely associated with a distribution shift by a portion of the NARW population to the Gulf of St. Lawrence in Canada during summer in the northern hemisphere (Meyer-Gutbrod et al., 2021), though the exact locations of individual entanglement events are largely equivocal despite increased monitoring since 2017. Long term declines in body size (Stewart et al., 2021) and overall health (Pirotta et al., 2023) could be indicative of trends in individual tendency toward the severity of entanglement events more so than a regime shift toward greater entanglement risk.

By assuming the absence of natural mortality, our model apportioned all mortality events between the anthropogenic causes of entanglement and vessel strike. Our justification was both empirical and theoretical: no necropsies of NARW carcasses (beyond the calf ages) have yielded a natural cause of death (Moore et al., 2004; Sharp et al., 2019) and natural mortality events are generally not expected for large long-lived mammals until senescence occurs close to the maximum lifespan (Anderson, 2018). Necropsies that have yielded an unknown cause of death typically occur due to the poor condition of the carcass (though not so poor as to obscure individual identification) or circumstances where the evidence is consistent with both anthropogenic causes (Moore et al., 2004; Sharp et al., 2019). As for natural mortality, while there is evidence that neurotoxins from harmful algal blooms (HABs) have been detected in NARWs and are associated with calf mortality events in southern right whales (*Eubalaena australis*) (see references in Moore et al. (2021)), there remains no evidence that HABs are responsible for adult NARW mortality. For practical reasons, a model that attempted to estimate a natural mortality rate was not identifiable with a near 100% overlap between prior and posterior distributions (not shown), which should be expected given the lack of observed data. Our model could be expanded to accommodate observations of natural or other mortality if they occur, though the likely sparse data would require careful modeling assumptions to facilitate estimation of any low hazard rates.

The poor identifiability of the recovery probability for vessel strike deaths warrants caution when interpreting the model results. The difficulty in estimation is likely due to the sparse observations of vessel strike severe injuries, which are limited to sharp traumas since blunt traumas can only be detected during necropsy (Sharp et al., 2019). While our random effects structure should add some robustness to the parameter estimation (Nater et al., 2020), it is clear that uncertainty in vessel strike carcass recovery induces uncertainty in the hazard rates (Figure S1) and ultimately the apportionment of cause to mortality (Figure 4). Higher probabilities of vessel strike carcass recovery suggest increased hazard rates of injury and mortality due to entanglement, acknowledging that a high recovery probability for vessel strikes likely indicates more unobserved deaths are due to entanglements. With lower recovery probabilities for vessel strike deaths, the corresponding hazard rates of injury and mortality for vessel strike are higher (and for entanglements, lower). Preliminary model fitting that separated sharp and blunt traumas as unique vessel strike events indicated additional identifiability problems, given that blunt traumas could not be observed as severe injuries.

Our effort here is one of numerous examples to estimate rangewide NARW survival (e.g., Schick et al. (2013); Pace et al. (2017); Reed et al. (2022); Pirotta et al. (2023)), with a variety of data constructs and model formulations. The rich sightings data that result from intensive survey efforts by the NARWC and photo analysis by the New England Aquarium enable many potential approaches to address different specific objectives. Our objective was to partition mortality between the anthropogenic causes hypothesized to be primarily responsible for the recent species decline to help facilitate management actions. The multistate injury and mortality model presented here could be further developed to be spatially explicit (e.g., Gowan et al. (2019)) or to accommodate smaller time steps (e.g., 3 months; Schick et al. (2013)); a continuous-time formulation (Rushing, 2023) might provide the greatest opportunity to address nuances in the hazard rates across space and time. An improved understanding of spatial and seasonal variation in NARW mortality risk across the western Atlantic may ultimately be necessary for effective conservation efforts to prevent further population decline and promote species recovery.

## Acknowledgements

We are grateful to the North Atlantic Right Whale Consortium (NARWC) for access to the sightings data. The capacity to develop precise estimates of North Atlantic right whale demographic parameters is due to the thousands of photographic captures of whales contributed by hundreds of collaborators working through the NARWC for nearly 40 years. We thank W. Getz, T. Branch, A. Read, and E. Cooch for helpful comments on an earlier version of this paper. Any use of trade, firm, or product names is for descriptive purposes only and does not imply endorsement by the U.S. Government.

## Supplementary Material

**Table S1:** Parameter estimates from multistate capture-recapture model of North Atlantic right whales during 1990–2019. Posterior summaries include mean, standard deviation (SD), quantiles describing the median and lower/upper 95% intervals, and the probability of inclusion.

| parameter | mean | SD | 2.5% | 50% | 97.5% | Pr(inclusion) |
| --- | --- | --- | --- | --- | --- | --- |
| mu.iE | -3.578 | 0.176 | -3.956 | -3.563 | -3.281 | 1 |
| mu.iV | -4.478 | 0.416 | -5.357 | -4.445 | -3.779 | 1 |
| mu.mE | -0.150 | 0.157 | -0.499 | -0.133 | 0.107 | 1 |
| mu.mV | 0.935 | 0.201 | 0.515 | 0.944 | 1.304 | 1 |
| beta.iE.FC | 1.082 | 0.467 | 0.000 | 1.180 | 1.750 | 0.90 |
| beta.iE.FR | -0.028 | 0.594 | -1.343 | 0.000 | 0.901 | 0.23 |
| beta.iE.age | -0.070 | 0.101 | -0.294 | 0.000 | 0.000 | 0.37 |
| beta.iE.regime | 0.603 | 0.106 | 0.400 | 0.601 | 0.820 | 1 |
| beta.iV.FC | -0.546 | 1.659 | -5.239 | 0.000 | 1.598 | 0.48 |
| beta.iV.FR | 0.739 | 0.687 | 0.000 | 0.795 | 1.988 | 0.66 |
| beta.iV.age | -0.112 | 0.151 | -0.438 | 0.000 | 0.000 | 0.41 |
| beta.iV.regime | 0.275 | 0.215 | -0.191 | 0.290 | 0.664 | 1 |
| beta.mE.FC | -0.042 | 0.525 | -1.086 | 0.000 | 0.573 | 0.18 |
| beta.mE.FR | 0.034 | 0.231 | -0.333 | 0.000 | 0.737 | 0.17 |
| beta.mE.age | -0.001 | 0.016 | 0.000 | 0.000 | 0.000 | 0.03 |
| beta.mV.FC | 0.468 | 1.923 | -3.642 | 0.000 | 5.358 | 0.46 |
| beta.mV.FR | 1.033 | 1.715 | -0.169 | 0.000 | 5.789 | 0.46 |
| beta.mV.age | 0.033 | 0.091 | 0.000 | 0.000 | 0.329 | 0.16 |
| psi.iE2 | 0.090 | 0.048 | 0.020 | 0.083 | 0.202 | 1 |
| mu.p | 1.351 | 0.044 | 1.267 | 1.350 | 1.438 | 1 |
| beta.p.sex | -0.196 | 0.058 | -0.310 | -0.196 | -0.081 | 1 |
| rho.inj | 0.923 | 0.030 | 0.853 | 0.927 | 0.971 | 1 |
| rho.mort | 0.687 | 0.057 | 0.571 | 0.689 | 0.792 | 1 |
| mu.rE | -1.897 | 0.307 | -2.464 | -1.911 | -1.257 | 1 |
| mu.rV | -0.829 | 0.546 | -1.698 | -0.894 | 0.408 | 1 |
| sigma.iE.t | 0.155 | 0.116 | 0.009 | 0.133 | 0.423 | 1 |
| sigma.iV.t | 0.384 | 0.177 | 0.095 | 0.370 | 0.760 | 1 |
| sigma.p.ind | 1.154 | 0.056 | 1.048 | 1.153 | 1.265 | 1 |
| sigma.p.t | 0.794 | 0.116 | 0.604 | 0.782 | 1.056 | 1 |
| sigma.r.t | 0.157 | 0.117 | 0.018 | 0.127 | 0.440 | 1 |
| delta | 0.459 | 0.019 | 0.421 | 0.459 | 0.497 | 1 |
| pi_age_known[1] | 0.576 | 0.020 | 0.537 | 0.576 | 0.615 | 1 |
| pi_age_known[2] | 0.032 | 0.007 | 0.020 | 0.032 | 0.048 | 1 |
| pi_age_known[3] | 0.019 | 0.006 | 0.010 | 0.019 | 0.031 | 1 |
| pi_age_known[4] | 0.019 | 0.006 | 0.010 | 0.019 | 0.032 | 1 |
| pi_age_known[5] | 0.353 | 0.019 | 0.315 | 0.353 | 0.391 | 1 |
| pi_age_unk[1] | 0.000 | 0.000 | 0.000 | 0.000 | 0.000 | 1 |
| pi_age_unk[2] | 0.536 | 0.090 | 0.359 | 0.537 | 0.708 | 1 |
| pi_age_unk[3] | 0.267 | 0.080 | 0.128 | 0.262 | 0.437 | 1 |
| pi_age_unk[4] | 0.131 | 0.060 | 0.039 | 0.123 | 0.270 | 1 |
| pi_age_unk[5] | 0.067 | 0.046 | 0.008 | 0.056 | 0.181 | 1 |

**Table S2:** Percentages of prior-posterior overlap for main parameters. While *μ* values were link-scale intercepts, all priors were specified on the real scale of the parameter and required transformation.

| Parameter | Prior | PPO% |
| --- | --- | --- |
| Mu.iE | Gamma(1,1) | 3 |
| Mu.iV | Gamma(1,1) | 2 |
| Mu.mE | Gamma(1,1) | 29 |
| Mu.mV | Gamma(1,1) | 21 |
| rho.inj | Uniform(0,1) | 15 |
| rho.mort | Uniform(0,1) | 28 |
| psi.iE2 | Uniform(0,1) | 22 |
| delta | Uniform(0,1) | 12 |
| Mu.p | Uniform(0,1) | 5 |
| Mu.rE | Beta(2,4) | 30 |
| Mu.rV | Beta(2,4) | 73 |

**Figure S1:**
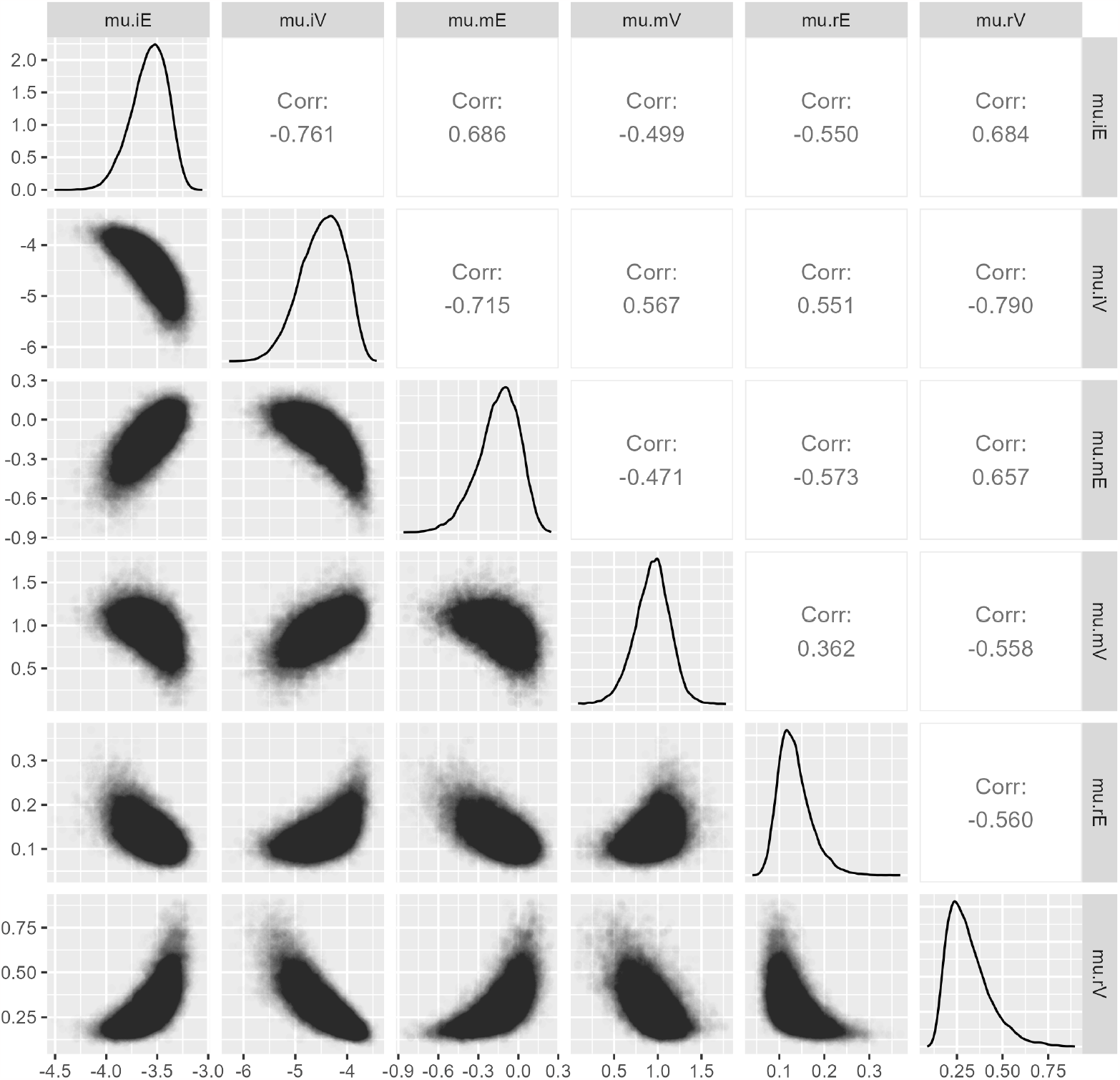
Posterior distributions and pairwise correlations of the link-scale intercepts for cause-specific (E = entanglement; V = vessel strike) injury (i) and mortality (m) rates, and recovery (r) probabilities. Rates are on the log scale while recovery probabilities are on the logit scale.

**Figure S2:**
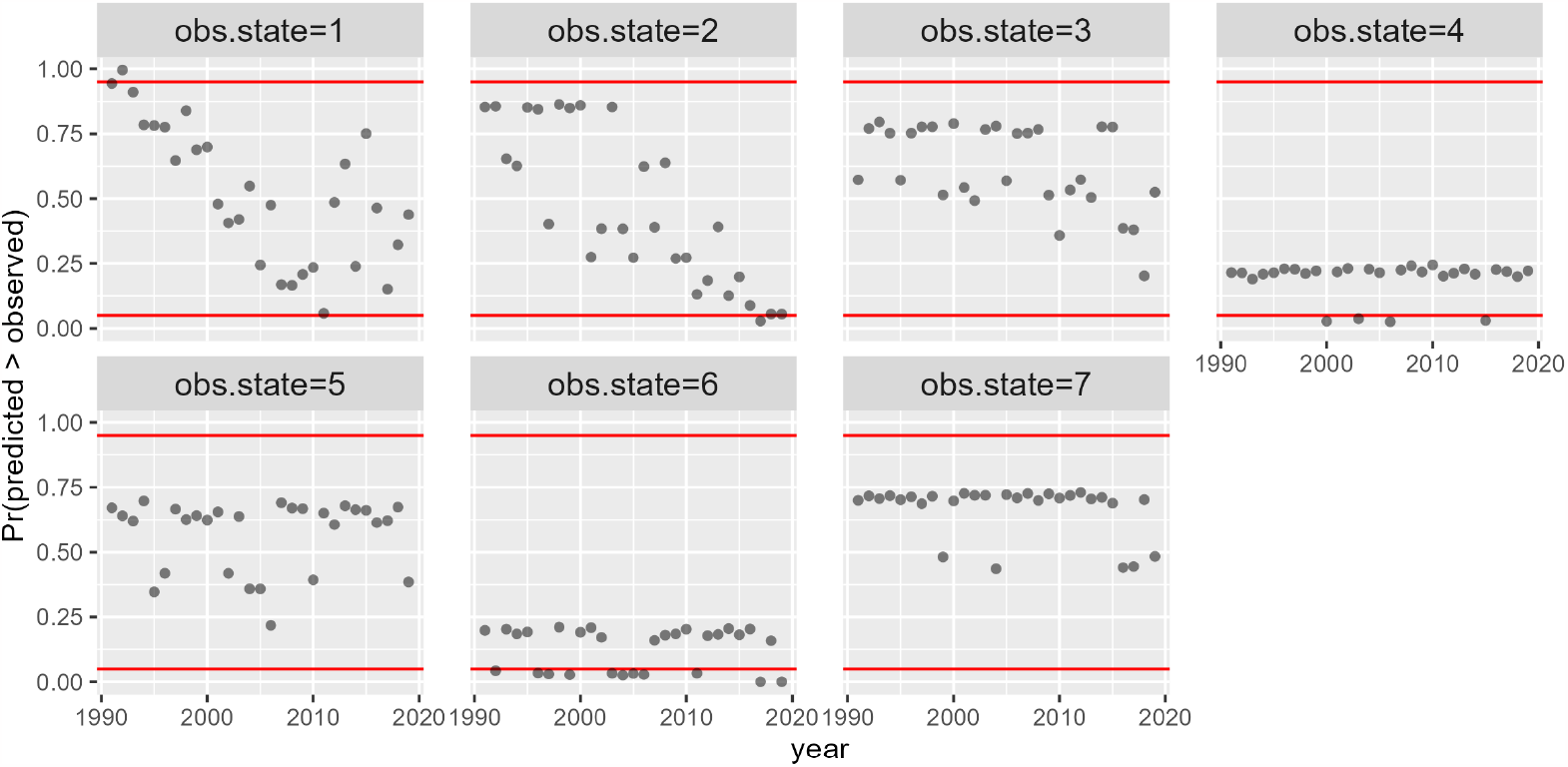
Posterior predictive check illustrating the proportion of simulated data sets that had a predicted count of events greater than the observed count of events for each event type (e.g., observation state) and year. Red lines indicate upper and lower thresholds (0.95 and 0.05, respectively) between which goodness-of-fit is adequate. Recorded events include: 1 = seen alive; 2 = seen injured by entanglement; 3 = recovered dead by entanglement; 4 = seen injured by vessel strike; 5 = recovered dead by vessel strike; 6 = seen injured by unknown cause; 7 = recovered dead by unknown cause.

## Footnotes

3 https://www.fisheries.noaa.gov/new-england-mid-atlantic/marine-life-distress/greater-atlantic-marine-mammal-stranding-network

4 https://www.dfo-mpo.gc.ca/species-especes/mammals-mammiferes/report-rapport/page01-eng.html

## References

Anderson, J.J., 2018. The relationship of mammal survivorship and body mass modeled by metabolic and vitality theories. Population Ecology 60, 111–125.

Benson, J.F., Dougherty, K.D., Beier, P., Boyce, W.M., Cristescu, B., Gammons, D.J., Garcelon, D.K., Higley, J.M., Martins, Q.E., Nisi, A.C., et al., 2023. The ecology of human-caused mortality for a protected large carnivore. Proceedings of the National Academy of Sciences 120, e2220030120.

Brooks, S., Gelman, A., 1998. General methods for monitoring convergence of iterative simulations. Journal of Computational and Graphical Statistics 7, 434–455.

Brownie, C., 1978. Statistical inference from band recovery data: a handbook. volume 131. US Department of the Interior, Fish and Wildlife Service.

Conn, P.B., Johnson, D.S., Williams, P.J., Melin, S.R., Hooten, M.B., 2018. A guide to bayesian model checking for ecologists. Ecological Monographs 88, 526–542.

Cristescu, B., Elbroch, L.M., Forrester, T.D., Allen, M.L., Spitz, D.B., Wilmers, C.C., Wittmer, H.U., 2022. Standardizing protocols for determining the cause of mortality in wildlife studies. Ecology and Evolution 12, e9034.

Davies, K.T., Brillant, S.W., 2019. Mass human-caused mortality spurs federal action to protect endangered north atlantic right whales in canada. Marine Policy 104, 157–162.

Davis, G.E., Baumgartner, M.F., Bonnell, J.M., Bell, J., Berchok, C., Bort Thornton, J., Brault, S., Buchanan, G., Charif, R.A., Cholewiak, D., et al., 2017. Long-term passive acoustic recordings track the changing distribution of north atlantic right whales (eubalaena glacialis) from 2004 to 2014. Scientific Reports 7, 13460.

de Valpine, P., Paciorek, C., Turek, D., Michaud, N., Anderson-Bergman, C., Obermeyer, F., Wehrhahn Cortes, C., Rodrìguez, A., Temple Lang, D., Paganin, S., 2022. NIMBLE: MCMC, Particle Filtering, and Programmable Hierarchical Modeling. URL: https://cran.r-project.org/package=nimble, doi:10.5281/zenodo.1211190. R package version 0.12.2.

de Valpine, P., Turek, D., Paciorek, C., Anderson-Bergman, C., Temple Lang, D., Bodik, R., 2017. Programming with models: writing statistical algorithms for general model structures with NIMBLE. Journal of Computational and Graphical Statistics 26, 403–413. doi:10.1080/10618600.2016.1172487.

Ergon, T., Borgan, Ø., Nater, C.R., Vindenes, Y., 2018. The utility of mortality hazard rates in population analyses. Methods in Ecology and Evolution 9, 2046–2056.

Gimenez, O., Morgan, B.J., Brooks, S.P., 2009. Weak identifiability in models for mark-recapture-recovery data, in: Thomson, D., Cooch, E., Conroy, M. (Eds.), Modeling demographic processes in marked populations. New York: Springer, pp. 1055–1067.

Goldstein, B., Turek, D., Ponisio, L., 2021. nimbleEcology: Distributions for Ecological Models in NIMBLE. URL: https://cran.r-project.org/package=nimbleEcology. R package version 0.4.1.

Gowan, T.A., Ortega-Ortiz, J.G., Hostetler, J.A., Hamilton, P.K., Knowlton, A.R., Jackson, K.A., George, R.C., Taylor, C.R., Naessig, P.J., 2019. Temporal and demographic variation in partial migration of the north atlantic right whale. Scientific Reports 9, 353.

Hamilton, P., Knowlton, A., Marx, M., 2007. Right whales tell their own stories: the photo-identification catalog. The urban whale: North Atlantic right whales at the crossroads. Harvard University Press, Cambridge, MA, 75–104.

Hayes, S.H., Josephson, E., Maze-Foley, K., Rosel, P.E., Wallace, J., 2022. U.S. Atlantic and Gulf of Mexico Marine Mammal Stock Assessments 2021. NOAA Tech Memo NMFS-NE-288. doi:10.25923/6tt7-kc16.

Heisey, D.M., Patterson, B.R., 2006. A review of methods to estimate cause-specific mortality in presence of competing risks. The Journal of Wildlife Management 70, 1544–1555.

Henry, A., Garron, M., Morin, D., Smith, A., Reid, A., Ledwell, W., Cole, T., 2021. Serious injury and mortality determinations for baleen whale stocks along the gulf of mexico, united states east coast, and atlantic canadian provinces, 2014-2018. US Dept Commer, Northeast Fisheries Science Center Reference Document 21–07 .

Knowlton, A., Robbins, J., Landry, S., McKenna, H., Kraus, S., Werner, T., 2016. Effects of fishing rope strength on the severity of large whale entanglements. Conservation Biology 30, 318–328.

Knowlton, A.R., Clark, J.S., Hamilton, P.K., Kraus, S.D., Pettis, H.M., Rolland, R.M., Schick, R.S., 2022. Fishing gear entanglement threatens recovery of critically endangered north atlantic right whales. Conservation Science and Practice 4, e12736.

Koons, D.N., Rockwell, R.F., Aubry, L.M., 2014. Effects of exploitation on an overabundant species: the lesser snow goose predicament. Journal of Animal Ecology 83, 365–374.

Kraus, S.D., Rolland, R.M., 2007. The Urban Whale: North Atlantic Right Whales at the Crossroads. Harvard University Press.

Kraus, S.D., Rolland, R.M., 2009. The urban whale: North Atlantic right whales at the crossroads. Harvard University Press.

Lebreton, J., Nichols, J., Barker, R., Pradel, R., Spendelow, J., 2009. Modeling individual animal histories with multistate capture–recapture models. Advances in Ecological Research 41, 87–173.

Lebreton, J.D., Burnham, K.P., Clobert, J., Anderson, D.R., 1992. Modeling survival and testing biological hypotheses using marked animals: a unified approach with case studies. Ecological Monographs 62, 67–118.

May, R., Reitan, O., Bevanger, K., Lorentsen, S.H., Nygård, T., 2015. Mitigating wind-turbine induced avian mortality: Sensory, aerodynamic and cognitive constraints and options. Renewable and Sustainable Energy Reviews 42, 170–181.

Mayne, B., Berry, O., Davies, C., Farley, J., Jarman, S., 2019. A genomic predictor of lifespan in vertebrates. Scientific Reports 9, 1–10.

Meyer-Gutbrod, E.L., Greene, C.H., Davies, K.T., 2018. Marine species range shifts necessitate advanced policy planning: The case of the north atlantic right whale. Oceanography 31, 19–23.

Meyer-Gutbrod, E.L., Greene, C.H., Davies, K.T., Johns, D.G., 2021. Ocean regime shift is driving collapse of the north atlantic right whale population. Oceanography 34, 22–31.

Moore, M., Knowlton, A., Kraus, S., McLellan, W., Bonde, R., 2004. Morphometry, gross morphology and available histopathology in north atlantic right whale (eubalaena glacialis) mortalities (1970-2002). Journal of Cetacean Research and Management 6, 199–214.

Moore, M.J., Mitchell, G.H., Rowles, T.K., Early, G., 2020. Dead cetacean? beach, bloat, float, sink. Frontiers in Marine Science 7, 333.

Moore, M.J., Rowles, T.K., Fauquier, D.A., Baker, J.D., Biedron, I., Durban, J.W., Hamilton, P.K., Henry, A.G., Knowlton, A.R., McLellan, W.A., et al., 2021. Review assessing north atlantic right whale health: Threats, and development of tools critical for conservation of the species. Diseases of Aquatic Organisms 143, 205–226.

Nater, C., Vindenes, Y., Aass, P., Cole, D., Langangen, Ø., Moe, S.J., Rustadbakken, A., Turek, D., Vøllestad, L.A., Ergon, T., 2020. Size-and stage-dependence in cause-specific mortality of migratory brown trout. Journal of Animal Ecology 89, 2122–2133.

Nichols, J.D., Runge, M.C., Johnson, F.A., Williams, B.K., 2007. Adaptive harvest management of north american waterfowl populations: a brief history and future prospects. Journal of Ornithology 148, 343–349.

Ogle, K., Barber, J., 2020. Ensuring identifiability in hierarchical mixed effects bayesian models. Ecological Applications 30, e02159.

Pace, III, R.M., Corkeron, P.J., Kraus, S.D., 2017. State–space mark–recapture estimates reveal a recent decline in abundance of north atlantic right whales. Ecology and Evolution 7, 8730–8741.

Pace, III, R.M., Williams, R., Kraus, S.D., Knowlton, A.R., Pettis, H.M., 2021. Cryptic mortality of north atlantic right whales. Conservation Science and Practice 3, e346.

Pettis, H., Rolland, R., Hamilton, P., Knowlton, A., Burgess, E., Kraus, S., 2017. Body condition changes arising from natural factors and fishing gear entanglements in north atlantic right whales eubalaena glacialis. Endangered Species Research 32, 237–249.

Pirotta, E., Schick, R.S., Hamilton, P.K., Harris, C.M., Hewitt, J., Knowlton, A.R., Kraus, S.D., Meyer-Gutbrod, E., Moore, M.J., Pettis, H.M., et al., 2023. Estimating the effects of stressors on the health, survival and reproduction of a critically endangered, long-lived species. Oikos, e09801.

Pradel, R., 2005. Multievent: an extension of multistate capture–recapture models to uncertain states. Biometrics 61, 442–447.

R Core Team, 2022. R: A Language and Environment for Statistical Computing. R Foundation for Statistical Computing. Vienna, Austria. URL: https://www.R-project.org/.

Rankin, R., Nicholson, K., Allen, S., Krützen, M., Bejder, L., Pollock, K., 2016. A full-capture hierarchical bayesian model of pollock’s closed robust design and application to dolphins. Frontiers in Marine Science 3, 25.

Reed, J., New, L., Corkeron, P., Harcourt, R., 2022. Multi-event modeling of true reproductive states of individual female right whales provides new insights into their decline. Frontiers in Marine Science 9, 994481.

Rolland, R.M., Schick, R.S., Pettis, H.M., Knowlton, A.R., Hamilton, P.K., Clark, J.S., Kraus, S.D., 2016. Health of north atlantic right whales eubalaena glacialis over three decades: from individual health to demographic and population health trends. Marine Ecology Progress Series 542, 265–282.

Rushing, C.S., 2023. An ecologist’s introduction to continuous-time multi-state models for capture–recapture data. Journal of Animal Ecology 92, 936–944.

Schick, R.S., Kraus, S.D., Rolland, R.M., Knowlton, A.R., Hamilton, P.K., Pettis, H.M., Kenney, R.D., Clark, J.S., 2013. Using hierarchical bayes to understand movement, health, and survival in the endangered north atlantic right whale. PloS One 8, e64166.

Sharp, S.M., McLellan, W.A., Rotstein, D.S., Costidis, A.M., Barco, S.G., Durham, K., Pitchford, T.D., Jackson, K.A., Daoust, P.Y., Wimmer, T., et al., 2019. Gross and histopathologic diagnoses from north atlantic right whale eubalaena glacialis mortalities between 2003 and 2018. Diseases of Aquatic Organisms 135, 1–31.

Stewart, J.D., Durban, J.W., Knowlton, A.R., Lynn, M.S., Fearnbach, H., Barbaro, J., Perryman, W.L., Miller, C.A., Moore, M.J., 2021. Decreasing body lengths in north atlantic right whales. Current Biology 31, 3174–3179.

Tassistro, E., Bernasconi, D.P., Rebora, P., Valsecchi, M.G., Antolini, L., 2020. Modeling the hazard of transition into the absorbing state in the illness-death model. Biometrical Journal 62, 836–851.

Tavecchia, G., Adrover, J., Navarro, A.M., Pradel, R., 2012. Modelling mortality causes in longitudinal data in the presence of tag loss: application to raptor poisoning and electrocution. Journal of Applied Ecology 49, 297–305.

Turek, D., de Valpine, P., Paciorek, C.J., 2016. E?icient markov chain monte carlo sampling for hierarchical hidden markov models. Environmental and Ecological Statistics 23, 549–564.

Van Der Hoop, J.M., Moore, M.J., Barco, S.G., Cole, T.V., Daoust, P.Y., Henry, A.G., McAlpine, D.F., McLellan, W.A., Wimmer, T., Solow, A.R., 2013. Assessment of management to mitigate anthropogenic effects on large whales. Conservation Biology 27, 121–133.

Williams, B.K., Nichols, J.D., Conroy, M.J., 2002. Analysis and management of animal populations. Academic press, San Diego, CA.

Williams, R., Gero, S., Bejder, L., Calambokidis, J., Kraus, S.D., Lusseau, D., Read, A.J., Robbins, J., 2011. Underestimating the damage: interpreting cetacean carcass recoveries in the context of the deepwater horizon/bp incident. Conservation Letters 4, 228–233.

